# CD38 biallelic loss is a recurrent mechanism of resistance to anti-CD38 antibodies in multiple myeloma

**DOI:** 10.1101/2024.12.17.628799

**Authors:** Benjamin Diamond, Linda Baughn, Mansour Poorebrahim, Alexandra M. Poos, Holly Lee, Marcella Kaddoura, J. Erin Wiedmeier-Nutor, Michael Durante, Gregory Otteson, Dragan Jevremovic, Hongwei Tang, Stefan Fröhling, Marc A. Baertsch, Marios Papadimitriou, Bachisio Ziccheddu, Tomas Jelinek, Cendrine Lemoine, Alexey Rak, Damian J. Green, Ola Landgren, Paola Neri, Leif Bergsagel, Esteban Braggio, Shaji Kumar, Marc S. Raab, Rafael Fonseca, Nizar Bahlis, Niels Weinhold, Francesco Maura

## Abstract

Monoclonal antibodies targeting CD38 are a therapeutic mainstay in multiple myeloma (MM). While they have contributed to improved outcomes, most patients still experience disease relapse, and little is known about tumor-intrinsic mechanisms of resistance to these drugs. Antigen escape has been implicated as a mechanism of tumor cell evasion in immunotherapy. Yet, it is unknown whether MM cells can develop permanent resistance to anti-CD38 antibodies by acquiring genomic events leading to biallelic disruption of the *CD38* gene locus. Here, by using whole genome and whole exome sequencing data from 701 newly diagnosed patients, 67 patients at relapse with naivety to anti-CD38 antibodies, and 50 patients collected at relapse following anti-CD38 antibodies. We report a loss of CD38 in 20% (10/50) of patients post-CD38 therapy, three of which exhibited a loss of both copies. Two of these cases showed convergent evolution where distinct subclones independently acquired similar advantageous variants. Functional studies on missense mutations involved in biallelic CD38 events revealed that two variants, L153H and C275Y, decreased binding affinity and antibody-dependent cellular cytotoxicity of the commercial antibodies Daratumumab and Isatuximab. However, a third mutation, R140G, conferred selective resistance to Daratumumab, while retaining sensitivity to Isatuximab. Clinically, patients with MM are often rechallenged with CD38 antibodies following disease progression and these data support a role for next generation sequencing to guide treatment selection.

## INTRODUCTION

Monoclonal antibodies directed against CD38 (daratumumab; Dara, isatuximab; Isa) have changed the therapeutic landscape for multiple myeloma (MM) and are now routinely administered across most disease settings^1-8^. Despite their efficacy, patients almost inevitably develop resistance to anti-CD38 therapy which is associated with poor outcomes^9^. Mechanisms of resistance to anti-CD38 therapy are heterogenous, in part due to the multiple modes of action which include antibody-dependent cellular cytotoxicity and cellular phagocytosis, complement-dependent cytotoxicity and immunomodulatory effects^10,11^. Recent findings suggest that reduction of MM CD38 cell-surface expression (i.e., antigenic escape) contributes to resistance^12,13^. However, in many patients, this downregulation is temporary and CD38 protein expression returns^14,15^. Primary resistance to anti-CD38 antibodies has also been linked to genomic drivers euch as *XBP1* loss, presence of high APOBEC mutational burden and genomic complexity^16-18^.

For BCMA-targeted immunotherapies, whole genome sequencing (WGS) studies have shown that prolonged exposure to the selective pressures promotes the acquisition of permanent antigen loss or selection of subclones harboring such events^19,20^. However, such genomic investigations have never been performed on post-anti-CD38 relapse samples, limiting our understanding of tumor-intrinsic mechanisms of resistance. This knowledge gap represents a critical need as MM patients are routinely retreated with anti-CD38 directed therapies, for which we have two current options and more in development^21-23^. In fact, in contrast to epigenetic and transcriptional mechanisms of resistance, the existence and detection of genomic events leading to permanent loss of CD38 would impact our clinical decision in re-challenging patients with CD38-directed therapy.

A recent study leveraging SNP-arrays reported an excess of monoallelic losses of the *CD38* gene locus following exposure to anti-CD38 antibodies. However, these losses were heterogenous, including large and whole arm/chromosome events, and could not directly be linked to anti-CD38 antibody exposure. Furthermore, the limited SNP-array resolution did not allow the interrogation of single nucleotide variants (SNV), indels and structural variants (SV)^24^. To comprehensively characterize mechanisms of resistance to anti-CD38 antibodies, we analyzed genomic data from a cohort of 50 MM patients who experienced relapse after anti-CD38 antibody therapy. The prevalence of biallelic inactivation of *CD38* was 6%, including evidence of convergent evolution wherein competing subclones lose *CD38* through independent mechanisms. Further, monoallelic loss was found in an additional 14%. These findings are in contrast to WGS from 701 newly diagnosed (ND) MM from CoMMpass and 72 WGS from CD38-antibody naïve relapsed MM, where no biallelic events were observed. Through functional work and three-dimensional protein modelling, we revealed the existence of missense variants that subvert CD38 structure and differentially affect the binding and killing by Daratumumab and Isatuximab, offering biologic rationale for using genomic profiling to guide re-challenge with anti-CD38 antibodies in MM.

## METHODS

### Study Cohort

We interrogated WGS (60-100x) and Whole Exome Sequencing (WES) data from 50 patients (Mayo Clinic; n = 28, Calgary University; n = 22^19^) with relapse following anti-CD38 monoclonal antibodies without restriction to line of use. Two additional cases (total n=52) with known aberrations of CD38 protein expression were included from Heidelberg University but were excluded from prevalence analyses to avoid a selection bias. 701 WGS (from patients with concomitant RNA-seq data) were imported from the MMRF’s CoMMpass trial (NCT01454297IA13, dbGap: phs001323.v3.p1)^25^ and 67 WGS from relapsed/refractory MM (RRMM) patients naïve to anti-CD38 antibodies^26^ (EGAS00001006538, EGAS00001004363, EGAS00001004805 and EGAS00001005973) were also included as controls. This study, adhering to the Declaration of Helsinki, received approval from the respective Review Boards of the University of Calgary, Heidelberg University Medical Faculty, and Mayo Clinic in Rochester and in Arizona. Patient participation was contingent on written informed consent. The study involved a total of 825 MM patients. Fluorescence-activated cell sorting (FACS) and bulk RNAseq were used to validate the effect of genomic events on *CD38* expression. Missense mutations predicted to affect anti-CD38 antibody binding were validated with binding and killing assays in CD38-mutant K562 cells. Finally, the effects of observed *CD38* mutant variants on CD38 protein structure and epitope binding were modeled with PyMOL.

### Next Generation Sequencing

DNA from CD138+ MM cells underwent WGS or WES using protcols detailed below. For samples collected at the University of Calgary, the New York Genome Center (NYGC) performed sequencing, employing the TruSeq DNA Nano Library Preparation Kit. For samples collected at Mayo Clinic Rochester and Arizona, libraries were prepared using the modified NEB Ultra II (New England Biolabs, Ipswich, MA) and the Nextera Flex systems (Illumina, San Diego, CA) and sequenced at the Mayo Clinic Genome Analysis Core. For samples collected at Mayo Clinic Arizona, Novogene performed the sequencing following random shearing of the genomic DNA, end-repaired, A-tailed and ligated with Illumina adapters. Sequencing was done on Illumina Novaseq 6000 sequencer. The subsequent data analysis utilized a suite of different calling tools, such as Mutect2, Strelka and Lancet for SNVs and indels; GATK4 and ASCAT for copy number variants (CNV), as described previously^19^. SVs were called using SvABA, Manta, and DELLY^27-29^. For the samples from Heidelberg University, DNA of CD138+ cell fractions were isolated using the Allprep Kit (Qiagen). WGS libraries were prepared with the Illumina TruSeq Nano DNA kit and sequenced on Illumina NovaSeq6000 Paired-end 150 bp S4. Raw sequencing data was processed and aligned to human reference genome build 37 version hs37d5 using the DKFZ OTP WGS pipeline^30^. ASCAT or ACEseq was used for CNV and estimation of purity and ploidy. Indels were called by Platypus^31^. SNVs were determined using samtools mpileup (v1.2.166-3)^81^. For the SNVs additional filtering steps were applied, including blacklist filtering^32^, fpfilter (https://github.com/genome/fpfilter-tool) and removal of events located in regions coding for immunoglobulins. SVs were called using SOPHIA (https://github.com/DKFZ-ODCF/SophiaWorkflow). For samples from all sources, SVs were manually curated to define complex events (i.e., templated insertions, chromothripsis, and chromoplexy) as described previously^33^. To determine the tumor clonal architecture, we used DPClust (https://github.com/Wedge-lab/dpclust). Phylogenetic trees were resolved according to the Pigeonhole principle^34^. All analyses were performed in R (v4.3.1).

### Flow Cytometry

Flow cytometry was performed as previously described^35^. Both Mayo Clinic and Calgary University followed this protocol, or similar. Briefly, PCPRO testing was performed on BM aspirate samples using antibodies to CD19, CD38, CD45, CD138, and kappa and lambda light chains, and 4′,6-diamidino-2-phenylindole (DAPI). 5×10^5^ events were acquired per sample, where available, using BD FACSCanto II (BD Biosciences, Franklin Lakes, NJ) and the analysis was performed by Kaluza software (Beckman Coulter Life Sciences, Indianapolis, IN). Abnormal clonal PCs were identified using differential expression of surface antigens, kappa-to-lambda ratio, and DAPI staining. Specifically, the CD38-multi-epitope-FITC (CYT-38F2, Cytognos) polyclonal antibody was employed to target the entire transmembrane molecule to prevent interference by residual bound anti-CD38 therapeutic antibody. In some cases, prior to therapy, a monoclonal CD38 APC (BD Biosciences) antibody was utilized.

### CD38 Binding Assay

To assess the binding affinity of CD38 mutants to daratumumab (Dara) and isatuximab (Isa), CD38 variants were expressed in K562 cells. Wild-type and mutant CD38 sequences were cloned into the lentiviral expression vector pLX307 (Addgene #41392). Lentiviral particles were produced and used to transduce K562 cells, followed by puromycin selection (2 µg/mL) to enrich for transduced populations.

Surface expression of CD38 was confirmed using monoclonal (BioLegend, #356610) and polyclonal (Cell Signaling Technology, #14637S) anti-CD38 antibodies. For binding assays, Dara and Isa were used as primary antibodies to stain CD38-expressing K562 cells. Subsequently, an anti-human IgG1 secondary antibody (SouthernBiotech, #9054-09) was employed to detect Dara-and Isa-bound cells. Non-transduced K562 cells served as a negative control to define gating parameters. To minimize nonspecific binding, Fc gamma receptor blockade (BioLegend #422302) was performed before each staining.

### Cytotoxicity of Daratumumab and Isatuximab against K562^CD38 R140G^ cells

To assess the cytotoxic effects of Daratumumab (Dara) and Isatuximab (Isa) on CD38^R140G^ mutant cells, a co-culture experiment was conducted using K562^CD38 WT^ and K562^CD38 R140G^ cells and varying concentrations of Dara or Isa in the presence of peripheral blood mononuclear cells (PBMCs) at an effector-to-target ratio of 20:1 over three days. Initially, K562 cells were seeded in round-bottom 96-well plates at a density of 1×10^4^ cells per well. Then, different concentrations of Dara and Isa were prepared and added to the respective wells. The mixture of K562 cells and antibodies was incubated for 30 minutes at 37°C. Subsequently, 2×10^5^ PBMCs were added to each well, and the plates were incubated at 37°C for three days. On the day 2, the co-cultures were again gently mixed to increase cell-cell contacts. The cytotoxicity was evaluated using flow cytometry by staining the cells with CD38 antibody and quantification of residual tumor cells within gated live cells. To enable complement-dependent cytotoxicity (CDC) in addition to antibody-dependent cellular cytotoxicity (ADCC), the experiments were carried out in a culture medium containing 10% human serum instead of fetal bovine serum (FBS). Controls containing tumor cells and Dara/Isa (without PBMCs) were included to remove the masking effect of Dara/Isa on CD38 detection.

## RESULTS

### Prevalence of deletions at the CD38 locus

To determine the prevalence of *CD38* loss, we examined 50 WGS from MM that relapsed following CD38-directed therapy and compared them to 701 patients with newly diagnosed MM (NDMM) from the CoMMpass dataset (NCT01454297) with available WGS and RNA-seq. The total number of cases with *CD38* deletions or copy neutral-loss of heterozygosity (CN-LOH) in the post-CD38-antibody cohort was significantly higher than in CoMMpass NDMM [20% (10/50) vs 7.1% (50/701), Fisher test, p = 0.004]. In our second control cohort of 67 anti-CD38 antibody naïve RRMM patients, 7 losses of the *CD38* locus (10.5%) were seen, which did not differ significantly from the frequency in anti-CD38-exposed MM (Fisher test, p = 0.2) or NDMM (Fisher test, p = 0.3, **Fig 1A-C**). Most of the events were large [>5 Megabase (MB)]^36^ or whole arm/whole chromosomal deletions including 48 (6.8%) cases in the NDMM, 9 (18%) cases in the post-CD38-antibody cohort, and all 7 (10.5%) cases in the anti-CD38-naïve RRMM (**Supplemental Table 1**). Leveraging WGS resolution and integrating CNV, SV, and SNV/indels, biallelic loss of the *CD38* locus at relapse following anti-CD38 antibody treatment was observed in 3/50 cases (6%; **Fig 1D**). Two of these patients had evidence of convergent evolution, where more than one subclone with biallelic loss of CD38 was detected. In addition to the three patients with biallelic loss, 8 (16%) relapsed with monoallelic deletions on *CD38*, one of which was defined as focal (i.e. <5 Mb). In contrast, there were no biallelic events and only 2 SNV, one nonsense and one missense, among the 701 NDMM cases (Fisher test, compared to post-CD38-directed therapy, p<0.001; **Supplemental Table 2**). Similarly, there was only 1 missense mutation (1.4%) in *CD38* in the anti-CD38 naïve RRMM and it was not in combination with a CNV. Of the two additional RRMM patients, selected based on strong evidence of CD38 downregulation, one had a biallelic event while the other had a nonsynonimus mutations detected by targeted sequencing. Combining these two patients with the others, a total of 6 nonsynonymous mutations in *CD38* were detected (**Supplemental Table 3**). Below, we provide details of the cases with biallelic events involving CD38 and the functional impact of CD38 nonsynonymous mutations in patients treated with anti-CD38 monoclonal antibody therapy.

**Figure 1.**
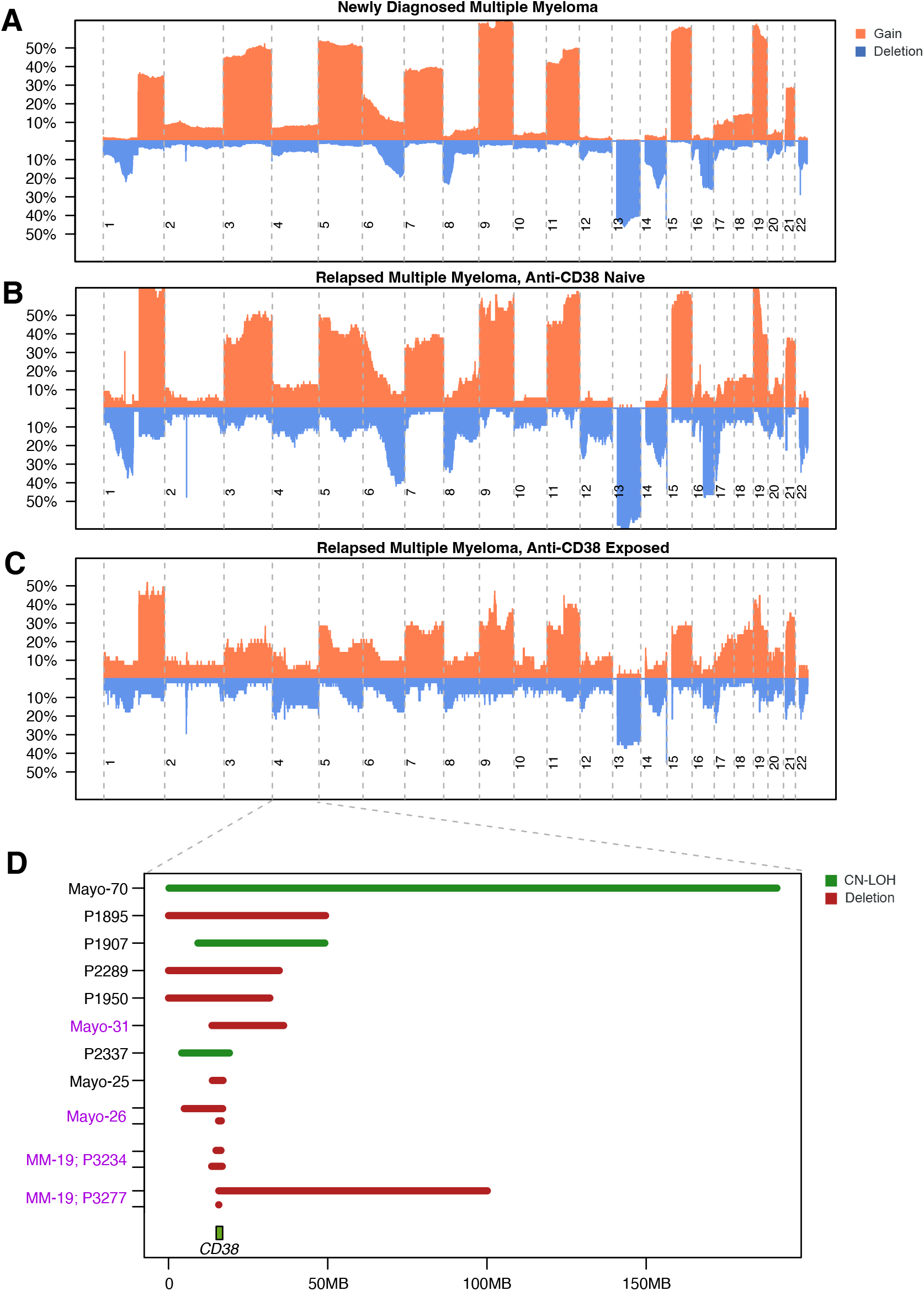
Loss of *CD38* is enriched in multiple myeloma relapsing after anti-CD38 antibodies. **A-**Cumulative copy number profiles of NDMM (**A**), anti-CD38-naïve RRMM (**B**), and anti-CD38-exposed RRMM (**C**); the latter with an increased frequency of losses involving the CD38 locus. Zoom-in plot of chromosome 4 showing summary of events involving the *CD38* locus. Samples with purple text are those identified as having biallelic events.

### Biallelic loss of CD38 and patterns of convergent evolution

In the first case, patient MM-19 received the triplet regimen Dara, bortezomib, and dexamethasone for 17 months with very good partial response (VGPR) as best response. Upon relapse, the patient received Dara in combination with a GPRC5D-directed bispecific antibody, resulting in a stringent complete response (CR) for 15 months at which time extramedullary myeloma was detected at multiple sites. Biopsies were obtained from two anatomically independent hepatic sites (**Fig 2A-B**) and WGS was performed^19^. A phylogenetic tree was constructed, classifying mutations as either truncal (i.e., shared and ancestral between both anatomic sites) or branched (i.e., unique to one sample; **Fig 2B**)^34,37,38^. In both disease sites, copy number loss resulted in biallelic deletion of the *CD38* locus. Demonstrating convergent evolution, the dominant subclone at each site had acquired a unique biallelic deletion. Specifically, one site (P3234) had two focal deletions encompassing the entire CD38 gene on chromosome 4, while the second site (P3277) displayed a focal deletion (∼27 kb) at the 5’ end of the CD38 gene along with a whole-arm loss of chromosome 4 (**Fig 2C**). Longitudinal flow cytometry analysis of bone marrow plasma cells at diagnosis and during treatment revealed CD38 expression at baseline, which was lost at relapse when WGS was performed. (**Fig 2D**).

**Figure 2.**
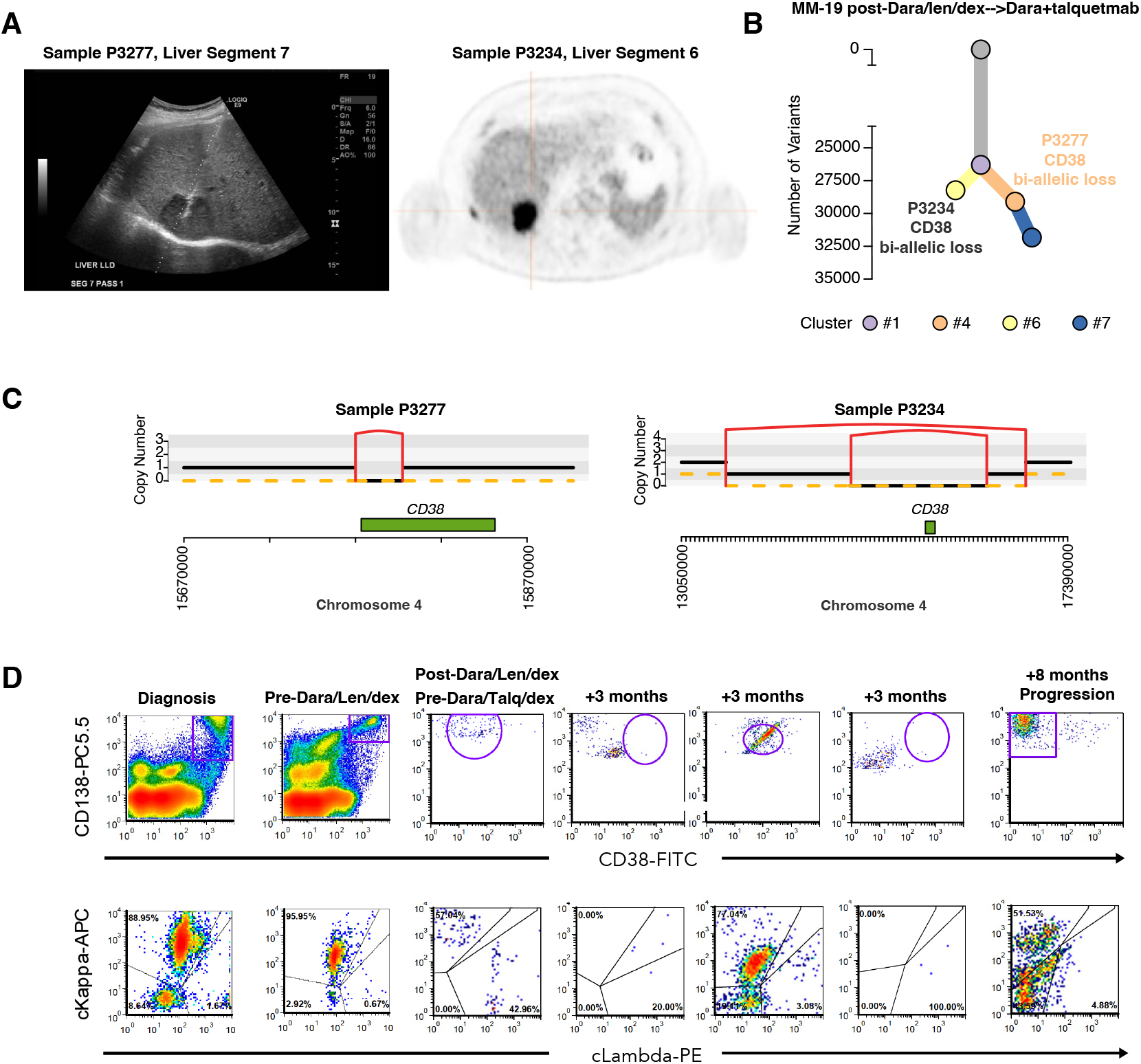
Convergent evolution towards biallelic loss of *CD38* at anatomically distinct sites of extramedullary disease. **A)** Ultrasound-guided biopsy of a hepatic segment 7 lesion (left) and positron-emission tomography-computed tomography of a hepatic segment 6 lesion (right). **B)** Phylogenetic tree reconstruction of WGS from both sites of disease. **C)** Mutographs of segments of chromosome 4 for each sample highlighting events involving the *CD38* locus. Horizontal black lines = total copy number, dashed lines = minor copy number, vertical red lines = deletions. **D)** Longitudinal FACS for patient MM-19 gated on CD38 and CD138 (top) with pull-down of selected events gated on kappa and lambda (bottom).

In the second case, patient Mayo-26 was treated with a seventh-line regimen of Dara/lenalidomide/dexamethasone, achieving a CR. However, relapse occured 15 months later. WGS revealed evidence of convergent evolution with two subclones with CD38 antigenic escape. The two subclones shared a common, large 12MB deletion on chromosome 4p but independently acquired a second hit in *CD38*. One subclone had loss of the second *CD38* allele via a reciprocal translocation involving the entire body of the gene at a cancer cell fraction (CCF) of 0.40. The other subclone harbored a missense mutation, resulting in the substitution of lysine for histidine at position 153 with a CCF of 0.60 (**Fig 3A-B**). Flow cytometric data at relapse revealed two CD138+ plasma cell populations: one lacked CD38 expression, consistent with biallelic deletion, while the other retained CD38 expression (i.e. L153H) (**Fig 3C**).

**Figure 3.**
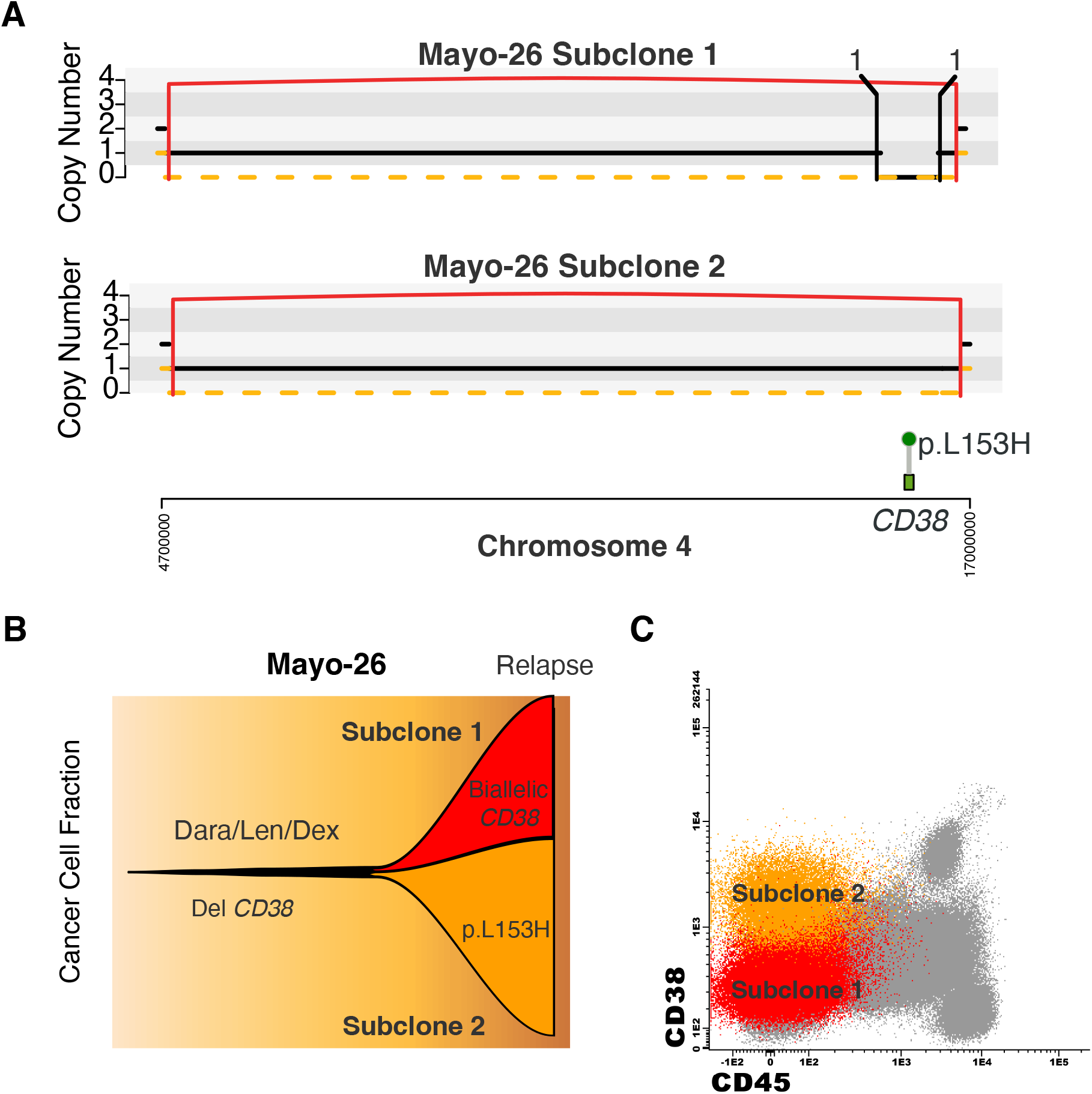
Biallelic events leading to divergent CD38 surface expression in distinct subclones. **A)** Two subclones sharing a common deletion on chromosome 4p encompassing *CD38*; one with balanced translocation disrupting the other allele and the other with a missense mutation at residue 153. **B)** Fish plot representation of CCFs. **C)** FACS from bone marrow at relapse showing two distinct myeloma populations, one with no CD38 expression corresponding to Subclone 1, and one with intact CD38 expression corresponding to Subclone 2.

In the third case, patient RRMM54 selected for sequencing because of a known decrease in CD38 expression, received five prior lines of therapy before receiving a combination of Dara, bortezomib and dexamethasone. At relapse, WGS revealed a clonal monoallelic deletion on 4 (**Fig 4A**). Two distinct variants were detected on the remaining CD38 allele, indicative of convergent evolution: 1) A single-nucleotide deletion of a cytosine base, resulting in a frameshift mutation at exon 2 (P98Lfs*12) with a CCF of 41%. A missense mutation substituting glycine for arginine at position 140 (R140G), observed at a higher CCF of 59% (**Fig. 4B**). Bulk RNA sequencing further revealed that CD38 expression in this patient was below the first decile compared to newly diagnosed MM samples in the MMRF CoMMpass dataset, in line with biallelic inactivation of the gene (**Fig 4B**).

**Figure 4.**
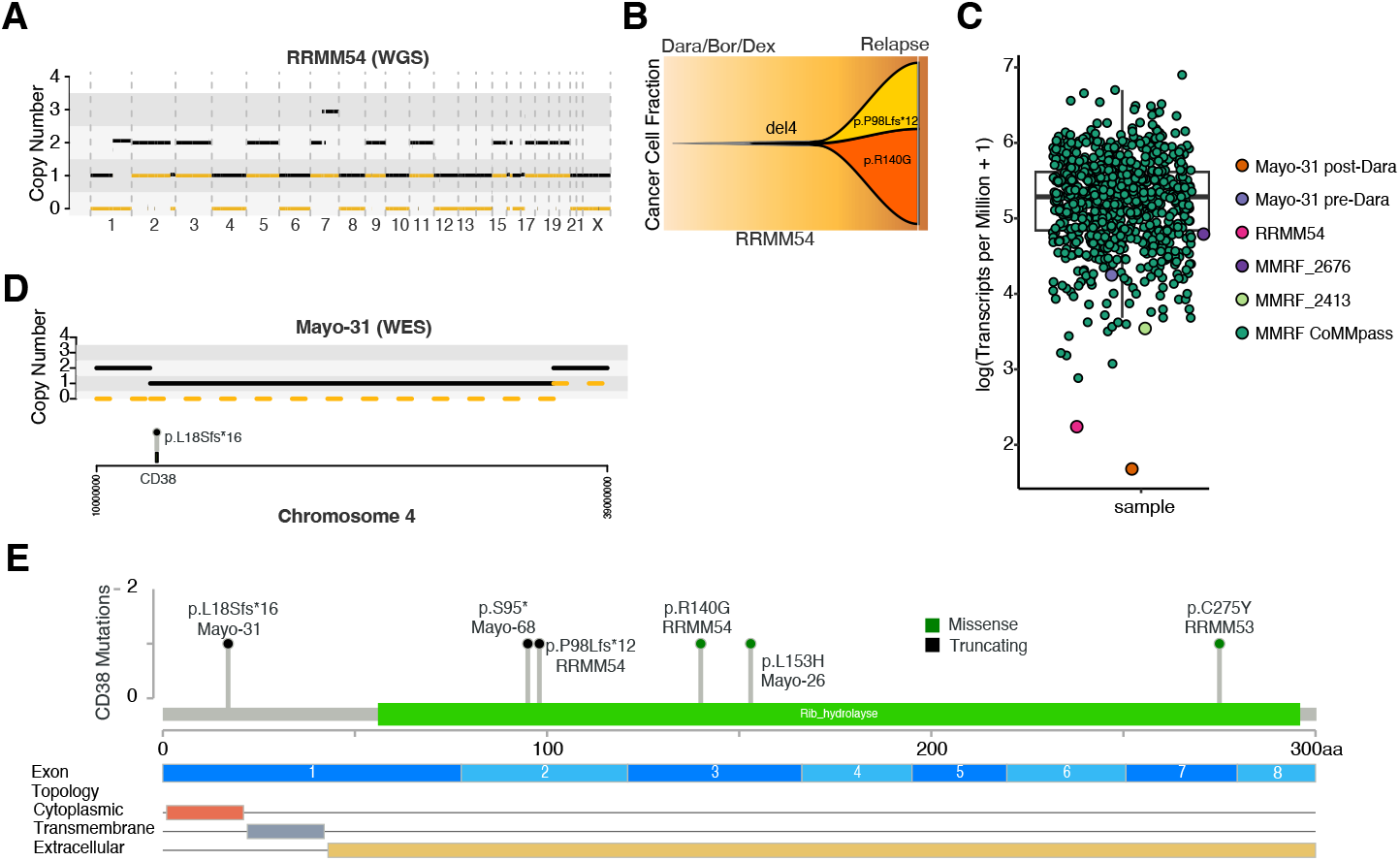
Biallelic events leading to decreased CD38 expression. **A)** Copy number profile from WGS for patient RRMM54 with loss of chromosome 4. **B)** FISH plot with CCFs of subclonal events on the remaining *CD38* allele with convergence towards *CD38* antigen escape. **C)** WES from patient Mayo-31 with biallelic inactivation of CD38. **D)** RNAseq CD38 expression from both antigen escape cases compared to baseline samples from CoMMpass. **E)** Summary of all *CD38* SNV and indels from anti-CD38 antibody relapses.

A fourth case, Mayo-31, had WES data from relapse after six months of Dara-based combination therapy revealing a 22MB deletion with a single nucleotide deletion of a guanine base in exon 1 (L18Sfs*16, CCF 62%) resulting in a frameshift deletion on the remaining allele associated with a 2.5-fold reduction in CD38 expression by RNAseq compared to baseline (**Fig 4C-D**).

Two other cases of note include a patient, Mayo-68, with a truncating mutation in a serine residue of exon 2 (S95*), seemingly a monoallelic event, at a CCF of 0.21, and a patient, RRMM53, with a missense mutation resulting in a substitution of Cysteine for Tyrosine at residue 275 (C275Y). The former case had sample collected after 30 months of intermittent Dara-based treatment and the latter case had sample collected after 17 months of Dara-based therapy (partial response) followed by 6 months of Isa-based therapy (minimal response). This C275Y mutation comes from targeted-sequencing in a low purity sample making it impossible to reliably reconstruct the copy number profile and status of the remaining allele. Finally, all mutations suspected to affect anti-CD38 antibody binding are summarized in **Fig 4E** and **Supplemental Table 2**.

### Impact of deletions on CD38 expression

As describe above, seven patients had monoallelic deletions or CN-LOH encompassing the *CD38* gene locus following CD38-directed therapy. One such patient, Mayo-25, had a focal monoallelic deletion of 3.3 MB. FACS from baseline revealed high CD38 expression, and the patient subsequently received 2 Dara-containing regimens. Ten months following the last Dara exposure, at relapse, FACS revealed vastly decreased CD38 expression suggesting that even acquired monoallelic events may play a role in resistance as has been previously reported (**Supplemental Fig 1**)^24,39^. Latency from last Dara exposure and the use of a polyclonal anti-CD38 antibody in the FACS protocol make interference from prior therapeutic antibody unlikely. Altogether, focal deletions (including cases with biallelic events) were enriched post-CD38-directed therapy (3/50, 6%) compared to NDMM (2/701, 0.3%; Fisher test, p = 0.002) and trended towards enrichment compared to CD38-naïve RRMM (0/67; Fisher test, p = 0.080). Interestingly, the 2 focal deletions in the CoMMpass NDMM were associated with low CD38 expression (i.e. in the first quartile) suggesting that focal events may indeed affect baseline resistance likely partnering with transcriptional suppression of the remaining allele (**Fig 4D**)^39^. Similarly, of the two SNV in *CD38* noted in baseline NDMM, MMRF_1796_1_BM had a monoallelic Q236* nonsense mutation and available RNAseq showed CD38 expression in the first quartile (**Supplemental Table 3**).

### Effects of CD38 variants on antibody function

To evaluate the functional impact of the variants identified in sequencing data, we transduced the K562 MM cell line to express the CD38 mutants S95*, L153H, C275Y, or R140G (**Methods**). Following puromycin selection of tranduced cells, we screened for surface protein expression using monoclonal and polyclonal CD38 antibodies (**Fig 5A**). Each variant except the truncated CD38 (S95*) was detectable, suggesting that this mutant is not trafficked to the cell surface. We next evaluated the binding affinity of Dara and Isa to the mutant CD38, revealing that neither commercial antibody bound to CD38 with L153H or C275Y. In contrast, both antibodies bound to R140G, calling into question its role as a resistance mechanism (**Fig 5B**). However, further binding assays for R140G at a range of concentrations revealed that Dara had a lower binding affinity for mutant protein compared to wild-type independent of drug concentration, while Isa showed similar binding to wild-type and R140G (**Fig 5C**). To determine whether the difference in binding activity translated into reduced therapeutic efficacy, we performed co-culture experiments with K562^CD38 WT^ and K562^CD38 R140G^ cells and varying concentrations of Dara or Isa in the presence of peripheral blood mononuclear cells (PBMCs). Diminished killing of K562^CD38 R140G^ cells was noted with Dara compared to Isa (**Fig 5D**).

**Figure 5.**
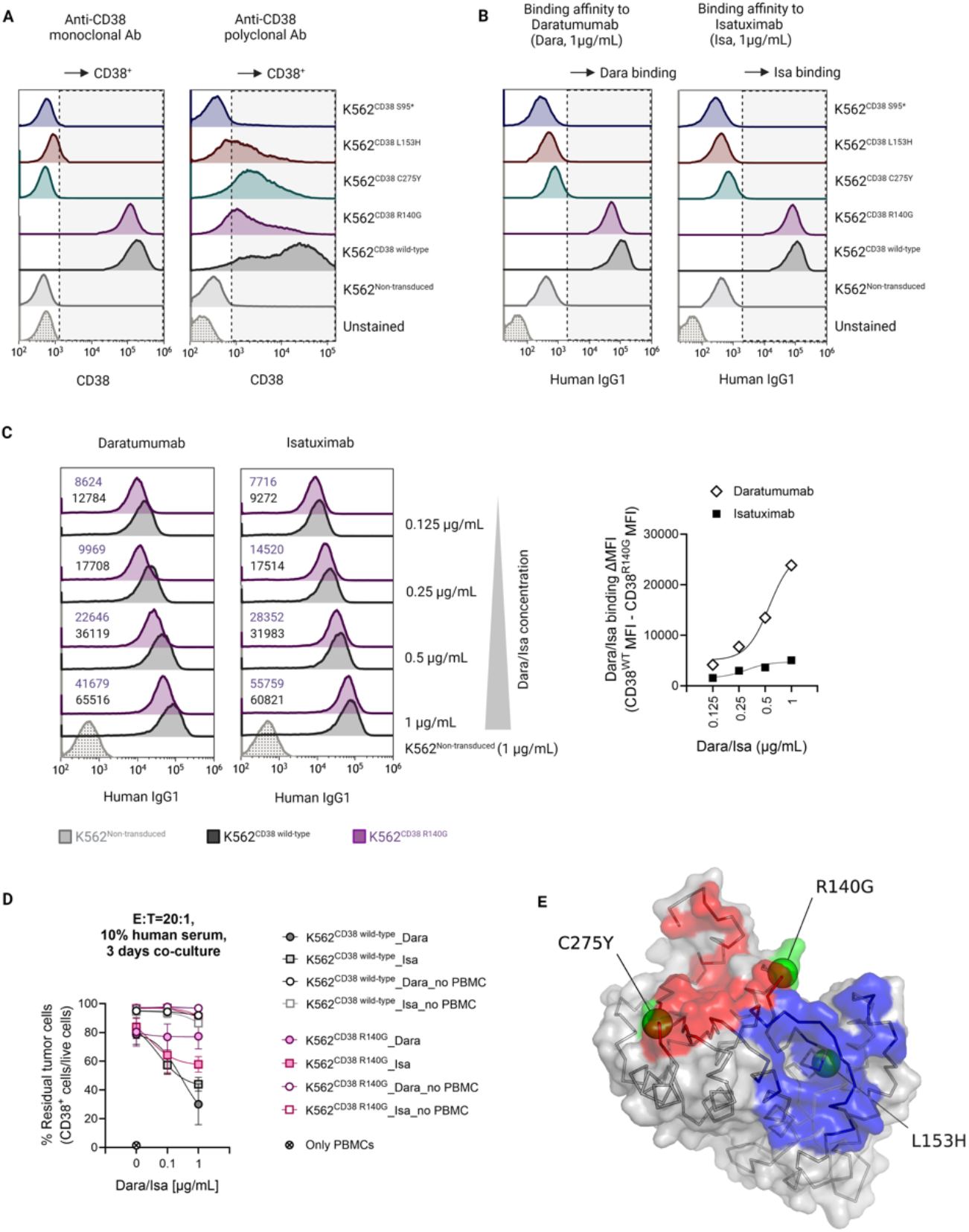
Functional validation of *CD38* variants. **A)** Wild-type and mutant CD38 protein expression in K562 cell lines by monoclonal and polyclonal anti-CD38 antibodies. **B)** CD38 binding assays: K562 cells were incubated with either Dara or Isa followed by secondary anti-IgG staining. **C)** Left: Repeated binding assay for K562^CD38 R140G^ (blue) at different drug concentrations with dose-independent decreased affinity for Dara compared to wild-type (grey). Right: Change in mean fluorescence intensity (MFI) recapitulating maintained binding of Isa for the R140G protein at all drug concentrations compared to the dose-independent decreased binding of Dara. **D)** Co-culture experiment with healthy donor peripheral blood mononuclear cells at Effector:Target ratio of 20:1 for various Dara and Isa concentrations demonstrating impaired antibody dependent cellular cytotoxicity of Dara compared to Isa for K562^CD38 R140G^ compared to K562^CD38 wild-type^. **E)** Structure of human CD38 (grey) with mapped epitopes for Isa (blue) and Dara (red), and three identified missense SNV (green spheres). Protein structure was visualized using PyMOL (The PyMOL Molecular Graphics System, Version 3.0, Schrödinger, LLC). The effect of R140G is described in text. Cys 275 forms a critical disulfide bond stabilising the protein structure. Mutation to bulky Tyr destabilizes CD38 structural integrity, affecting both epitopes. The aliphatic Leu 153 contributes to CD38 hydrophobic core formation under the Isa epitope. Its mutation to a charged His will destabilize the Isa epitope structural integrity affecting the antibody binding.

Finally, we modeled CD38 variants with respect to the binding epitopes of both commercial antibodies^40,41^. While C275Y and L153H both destabilize CD38 structural integrity, potentially affecting the binding site of both antibodies, Arginine 140 is located near the Dara binding site and mutation to a glycine increases flexibility of the neighboring amino acid residues forming the epitope and likely affects Dara binding (**Fig 5E**), but not Isa. Overall, this data reveals that mutations in *CD38* may still result in expressed protein, but selectively impair binding and efficacy of specific anti-CD38 therapies.

## DISCUSSION

Anti-CD38 antibodies have become the backbone of many MM combination therapies and virtually all patients will be exposed to this class, perhaps repeatedly, along their treatment course. Resistance to therapy will develop in the majority of patients and while loss of CD38 target antigen has been proposed as an etiology, biallelic inactivation of *CD38* has not been previously described^24^. Here, we observe that it is an important, yet relatively rare occurrence, likely responsible for resistance in 5-10% of cases. We reveal that CD38 expression may be lost to biallelic inactivation of the gene locus, via combinations of SV and truncating or nonsense mutations. We also srevealed that missense mutations may alter the structure of CD38 and disrupt binding epitopes of commercial antibodies without altering the CD38 surface expression. Importantly, resistance to one agent does not guarantee resistance to another. We demonstrated that an R140G mutation, acquired following Dara exposure, conferred resistance to Dara, but not Isa. Although rare, this data might justify the practice of substituting one drug for the other if anti-CD38 retreatment is attempted. Certain mutations produce expressed protein product that is resistant to both commercial agents. Notably, clinical FACS reveals cell surface expression of mutant CD38, which could lead to the futile use of anti-CD38 antibody rechallenge.

While genomic antigenic escape appears to be a relatively rare mechanism of resistance to anti-CD38 antibodies, it is striking that the majority of cases arose via convergent evolution; a signal of the profound selection pressure of prolonged antigen targeting^19,20^. In fact, all patients with convergent evolution were responders to anti-CD38 therapy, with at least 8 months of disease control. The demonstration that multiple subclones may independently converge on the same route of resistance also suggests that a degree of genomic instability may be required to facilitate this phenomenon, as has been seen in a recent large-scale classification study^36^. Apart from biallelic events, monoallelic loss of the *CD38* locus appears more frequently in post-CD38-antibody relapse samples as compared to baseline. Similarly to what has been reported by Portuguese et al., the overall prevalence of *CD38* loss here was 20%^24^. Although the remaining allele appears intact, a fraction of these events—particularly the focal ones—can significantly downregulate CD38 expression and may serve as a mechanism of resistance. As practice patterns shift and anti-CD38 consolidation and maintenance strategies becomes common practice^1,3,5,6,42^, these genomic resistance mechanisms are likely to emerge more frequently and earlier in the disease course.

The resolution of WGS has offered insights into antigen escape mechanisms that emerge at relapse following immunotherapy. Recent studies have indicated that these mechanisms are far more common than previously thought^19^, and that antigen escape mutations are typically acquired during treatment rather than being present at diagnosis^20^. However, larger scale and uniform genomic studies of relapsed disease following anti-CD38 antibody therapy will be required to identify whether distinct patterns of genomic instability predispose tumors to antigen escape^18,36^. While more work will need to be done in this space, the acquisition and selection of these alterations further supports the need for development of genomics platforms for longitudinal disease monitoring and personalized strategies in MM.

## DATA SHARING STATEMENT

Data are available at the following public repositories. WGS and WES data from 50 patients (Mayo Clinic; n = 28, Calgary University; n = 22^19^) with relapse following anti-CD38. Two additional cases (total n=52) with known CD38 downregulation were included from Heidelberg University but were excluded from prevalence analyses (EGAS50000000709). 701 WGS from MMRF’s CoMMpass trial (NCT01454297IA13, dbGap: phs001323.v3.p1)^25^ and 67 WGS from relapsed/refractory cases from patients naïve to anti-CD38 antibodies^26^ (EGAS00001006538, EGAS00001004363, EGAS00001004805 and EGAS00001005973).

## ACKNOWLEDGEMENTS

This work was supported by the Myeloma Solutions Fund (MSF), Paula and Rodger Riney Multiple Myeloma Research Program Fund, the Tow Foundation, and the Sylvester Comprehensive Cancer Center NCI Core Grant (P30 CA 240139).

FM is supported by the American Society of Hematology (ASH), Leukemia & Lymphoma Society (LLS), NIH-NCI, and by International Myeloma Society (IMS).

B.D. is supported by the National Cancer Institute of the National Institutes of Health under Award Number K12CA226330.

The authors are grateful to the patients for participation in this study. The Heidelberg Team thanks the Sample Processing Lab of the National Center for Tumor Diseases (NCT) Heidelberg, the Next-Generation Sequencing and Omics IT and Data Management Core Facilities of the German Cancer Research Center (DKFZ), the Biobank Multiple Myeloma UKHD and the Myeloma Registry at UKHD for excellent services. Support and funding of the project via the NCT Heidelberg Molecular Precision Oncology Program (projects H021, H067 and K08K) and the Dietmar-Hopp Foundation is gratefully acknowledged. A.P. is funded by the Medical Data Scientist Program of Heidelberg University, Faculty of Medicine. Data storage service via SDS@hd is supported by the Ministry of Science, Research and the Arts Baden-Württemberg (MWK) and the German Research Foundation (DFG) through grants INST 35/1314-1 FUGG and INST 35/1503-1 FUGG.

## CONTRIBUTIONS

N.J.B., N.W., R.F., and F.M. conceived of, designed and supervised all experiments, performed the analysis, and wrote the paper. B.D., A.P., L.B., and M.Po. analyzed the data, performed analyses, and wrote the paper. P.N., S.F., E.B., S.K., M.R., and O.L. contributed to patient consent and sample collection. M.K., J.E.W, M.D., G.O., D.G., D.J., H.T., M.B., M.Pa., B.Z., T.J., C.L, A.R., analyzed the data and performed analyses.

## REFERENCES

1. Facon T, Dimopoulos M-A, Leleu XP, et al. Isatuximab, Bortezomib, Lenalidomide, and Dexamethasone for Multiple Myeloma. New England Journal of Medicine. 2024.

2. Facon T, Kumar S, Plesner T, et al. Daratumumab plus lenalidomide and dexamethasone for untreated myeloma. New England Journal of Medicine. 2019;380(22):2104–2104.

3. Leleu X, Hulin C, Lambert J, et al. Isatuximab, lenalidomide, dexamethasone and bortezomib in transplant-ineligible multiple myeloma: the randomized phase 3 BENEFIT trial. Nature Medicine. 2024:1–7.

4. Leypoldt LB, Tichy D, Besemer B, et al. Isatuximab, carfilzomib, lenalidomide, and dexamethasone for the treatment of high-risk newly diagnosed multiple myeloma. Journal of Clinical Oncology. 2024;42(1):26–26.

5. Sonneveld P, Dimopoulos MA, Boccadoro M, et al. Daratumumab, bortezomib, lenalidomide, and dexamethasone for multiple myeloma. New England Journal of Medicine. 2024;390(4):301–301.

6. Voorhees PM, Kaufman JL, Laubach J, et al. Daratumumab, lenalidomide, bortezomib, and dexamethasone for transplant-eligible newly diagnosed multiple myeloma: the GRIFFIN trial. Blood, The Journal of the American Society of Hematology. 2020;136(8):936–936.

7. Attal M, Richardson PG, Rajkumar SV, et al. Isatuximab plus pomalidomide and low-dose dexamethasone versus pomalidomide and low-dose dexamethasone in patients with relapsed and refractory multiple myeloma (ICARIA-MM): a randomised, multicentre, open-label, phase 3 study. The Lancet. 2019;394(10214):2096–2096.

8. Dimopoulos MA, Oriol A, Nahi H, et al. Daratumumab, lenalidomide, and dexamethasone for multiple myeloma. New England Journal of Medicine. 2016;375(14):1319–1319.

9. Gandhi UH, Cornell RF, Lakshman A, et al. Outcomes of patients with multiple myeloma refractory to CD38-targeted monoclonal antibody therapy. Leukemia. 2019;33(9):2266–2266.

10. Saltarella I, Desantis V, Melaccio A, et al. Mechanisms of resistance to anti-CD38 daratumumab in multiple myeloma. Cells. 2020;9(1):167.

11. Van de Donk NW, Usmani SZ. CD38 antibodies in multiple myeloma: mechanisms of action and modes of resistance. Frontiers in immunology. 2018;9:2134.

12. Ise M, Matsubayashi K, Tsujimura H, Kumagai K. Loss of CD38 expression in relapsed refractory multiple myeloma. Clinical Lymphoma Myeloma and Leukemia. 2016;16(5):e59–e64.

13. Minarik J, Novak M, Flodr P, et al. CD 38-negative relapse in multiple myeloma after daratumumab-based chemotherapy. European journal of haematology. 2017;99(2):186–186.

14. Mikhael J, Belhadj-Merzoug K, Hulin C, et al. A phase 2 study of isatuximab monotherapy in patients with multiple myeloma who are refractory to daratumumab. Blood cancer journal. 2021;11(5):89.

15. Perez de Acha O, Reiman L, Jayabalan DS, et al. CD38 antibody re-treatment in daratumumab-refractory multiple myeloma after time on other therapies. Blood Advances. 2023;7(21):6430–6430.

16. Cohen YC, Zada M, Wang S-Y, et al. Identification of resistance pathways and therapeutic targets in relapsed multiple myeloma patients through single-cell sequencing. Nature medicine. 2021;27(3):491–491.

17. Maura F, Boyle EM, Coffey D, et al. Genomic and immune signatures predict clinical outcome in newly diagnosed multiple myeloma treated with immunotherapy regimens. Nature cancer. 2023;4(12):1660–1660.

18. Ziccheddu B, Giannotta C, D’Agostino M, et al. Genomic and immune determinants of resistance to daratumumab-based therapy in relapsed refractory multiple myeloma. Blood Cancer Journal. 2024;14(1):117.

19. Lee H, Ahn S, Maity R, et al. Mechanisms of antigen escape from BCMA-or GPRC5D-targeted immunotherapies in multiple myeloma. Nature Medicine. 2023:1–12.

20. Papadimitriou M, Ahn S, Diamond B, et al. Timing antigenic escape in multiple myeloma treated with T-cell redirecting immunotherapies. bioRxiv. 2024:2024.2005.2022.595383.

21. Grab AL, Kim PS, John L, et al. Pre-Clinical Assessment of SAR442257, a CD38/CD3xCD28 Trispecific T Cell Engager in Treatment of Relapsed/Refractory Multiple Myeloma. Cells. 2024;13(10):879.

22. Grandclément C, Estoppey C, Dheilly E, et al. Development of ISB 1442, a CD38 and CD47 bispecific biparatopic antibody innate cell modulator for the treatment of multiple myeloma. Nature Communications. 2024;15(1):2054.

23. Mei H, Li C, Jiang H, et al. A bispecific CAR-T cell therapy targeting BCMA and CD38 in relapsed or refractory multiple myeloma. Journal of Hematology & Oncology. 2021;14:1–17.

24. Portuguese AJ, Fang M, Tuazon SA, et al. Acquired CD38 gene deletion as a mechanism of tumor antigen escape in multiple myeloma. Blood Advances. 2023;7(23):7235–7235.

25. Skerget S, Penaherrera D, Chari A, et al. Comprehensive molecular profiling of multiple myeloma identifies refined copy number and expression subtypes. Nature Genetics. 2024:1–12.

26. Poos AM, Prokoph N, Przybilla MJ, et al. Resolving therapy resistance mechanisms in multiple myeloma by multiomics subclone analysis. Blood. 2023;142(19):1633–1633.

27. Rausch T, Zichner T, Schlattl A, Stütz AM, Benes V, Korbel JO. DELLY: structural variant discovery by integrated paired-end and split-read analysis. Bioinformatics. 2012;28(18):i333–i339.

28. Chen X, Schulz-Trieglaff O, Shaw R, et al. Manta: rapid detection of structural variants and indels for germline and cancer sequencing applications. Bioinformatics. 2016;32(8):1220–1222.

29. Wala JA, Bandopadhayay P, Greenwald NF, et al. SvABA: genome-wide detection of structural variants and indels by local assembly. Genome research. 2018;28(4):581–581.

30. Reisinger E, Genthner L, Kerssemakers J, et al. OTP: An automatized system for managing and processing NGS data. Journal of biotechnology. 2017;261:53–62.

31. Rimmer A, Phan H, Mathieson I, et al. Integrating mapping-, assembly-and haplotype-based approaches for calling variants in clinical sequencing applications. Nature genetics. 2014;46(8):912–912.

32. Danecek P, Bonfield JK, Liddle J, et al. Twelve years of SAMtools and BCFtools. Gigascience. 2021;10(2):giab008.

33. Rustad EH, Yellapantula VD, Glodzik D, et al. Revealing the impact of structural variants in multiple myeloma. Blood cancer discovery. 2020;1(3):258.

34. Nik-Zainal S, Van Loo P, Wedge DC, et al. The life history of 21 breast cancers. Cell. 2012;149(5):994–994.

35. Panakkal V, Lakshman A, Shi M, et al. Utility of flow cytometry screening before MRD testing in multiple myeloma. Blood cancer journal. 2023;13(1):55.

36. Maura F, Rajanna AR, Ziccheddu B, et al. Genomic classification and individualized prognosis in multiple myeloma. Journal of Clinical Oncology. 2024;42(11):1229–1229.

37. Maura F, Bolli N, Angelopoulos N, et al. Genomic landscape and chronological reconstruction of driver events in multiple myeloma. Nature communications. 2019;10(1):3835.

38. Landau HJ, Yellapantula V, Diamond BT, et al. Accelerated single cell seeding in relapsed multiple myeloma. Nature communications. 2020;11(1):3617.

39. Cirrincione AM, Poos AM, Ziccheddu B, et al. The biological and clinical impact of deletions before and after large chromosomal gains in multiple myeloma. Blood. 2024;144(7):771–771.

40. Lee HT, Kim Y, Park UB, Jeong TJ, Lee SH, Heo Y-S. Crystal structure of CD38 in complex with daratumumab, a first-in-class anti-CD38 antibody drug for treating multiple myeloma. Biochemical and Biophysical Research Communications. 2021;536:26–31.

41. Martin TG, Corzo K, Chiron M, et al. Therapeutic opportunities with pharmacological inhibition of CD38 with isatuximab. Cells. 2019;8(12):1522.

42. Badros AZ, Foster L, Anderson Jr LD, et al. Daratumumab with lenalidomide as maintenance after transplant in newly diagnosed multiple myeloma: the AURIGA study. Blood Journal. 2024:blood.2024025746.

