## Supplementary Figure for "CD38 biallelic loss is a recurrent mechanism of resistance to anti-CD38 antibodies in multiple myeloma"

**Supplemental Figure 1**: CD38 expression may be affected by monoallelic deletions. **A)** Baseline bone marrow FACS from Mayo-25 showing high CD38 expression. **B)** Relapse FACS at ten months following progressive disease to both Dara-Pomalidomide-Dexamethasone and Dara-Carfilzomib-Dexamethasone. **C)** Change in median fluorescence intensity (MFI) for CD38 across samples. **D)** Copy Number plot displaying the focal monoallelic deletion of *CD38*.

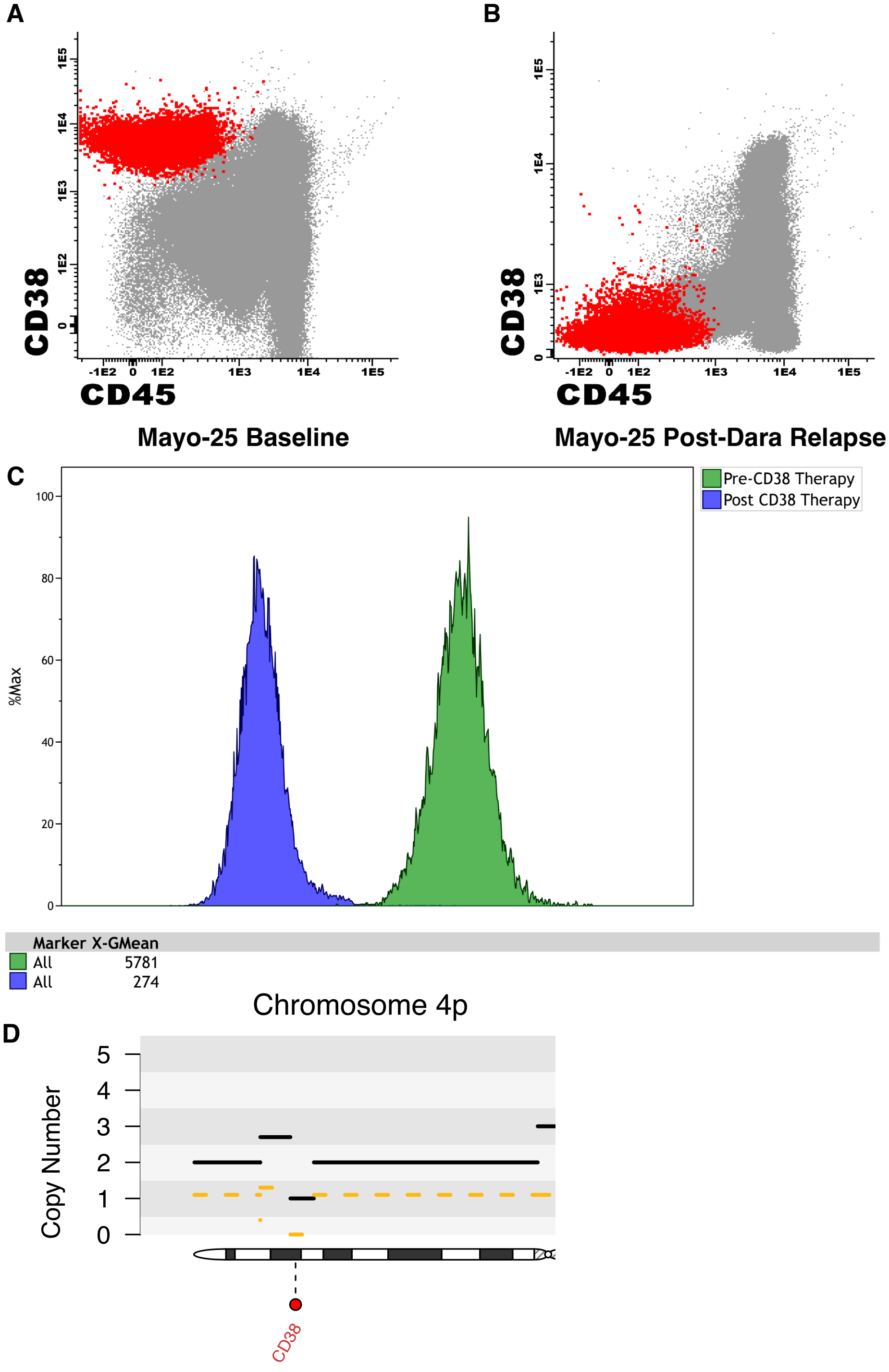
